# FoxMask: a new automated tool for animal detection in camera trap images

**DOI:** 10.1101/640037

**Authors:** Eric Devost, Sandra Lai, Nicolas Casajus, Dominique Berteaux

## Abstract

1. Camera traps now represent a reliable, efficient and cost-effective technique to monitor wildlife and collect biological data in the field. However, efficiently extracting information from the massive amount of images generated is often extremely time-consuming and may now represent the most rate-limiting step in camera trap studies.
2. To help overcome this challenge, we developed FoxMask, a new tool performing the automatic detection of animal presence in short sequences of camera trap images. FoxMask uses background estimation and foreground segmentation algorithms to detect the presence of moving objects (most likely, animals) on images.
3. We analyzed a sample dataset from camera traps used to monitor activity on arctic fox *Vulpes lagopus* dens to test the parameter settings and the performance of the algorithm. The shape and color of arctic foxes, their background at snowmelt and during the summer growing season were highly variable, thus offering challenging testing conditions. We compared the automated animal detection performed by FoxMask to a manual review of the image series.
4. The performance analysis indicated that the proportion of images correctly classified by FoxMask as containing an animal or not was very high (> 90%). FoxMask is thus highly efficient at reducing the workload by eliminating most false triggers (images without an animal). We provide parameter recommendations to facilitate usage and we present the cases where the algorithm performs less efficiently to stimulate further development.
5. FoxMask is an easy-to-use tool freely available to ecologists performing camera trap data extraction. By minimizing analytical time, computer-assisted image analysis will allow collection of increased sample sizes and testing of new biological questions.

## INTRODUCTION

Camera trapping is a very efficient and non-invasive method widely used by ecologists to collect biological data (Cutler & Swann 1999). In recent years, the use of camera traps has rapidly increased in several ecological field studies (Rowcliffe & Carbone 2008; McCallum 2013). Camera traps can be set to run continuously, to take images at specific time intervals (time-lapse setting) or to be motion-triggered (Cutler & Swann 1999; Meek & Pittet 2012). While the use of camera traps has many benefits, it can also generate large numbers of images (in the millions for many studies) in short periods of time, which may rapidly generate an overwhelming amount of data to treat. Several software are now available to facilitate data storage and management (e.g., Cpw Photo Warehouse, Ivan & Newkirk 2016; CamtrapR, Niedballa, Sollmann, Courtiol & Wilting 2016; ViXeN, Ramachandran, Devarajan & Poisot 2018), but efficiently extracting information from the images still remains a challenge. Going through images manually may now be the most rate-limiting step and often requires personnel specifically assigned to the task. When camera traps are in time-lapse setting, the number of uninformative images can be extremely high (Price Tack et al. 2016). Even when camera traps are motion-triggered, false triggers due to the wind, sun or moving vegetation increase the amount of images generated. To overcome some of these issues, image analysis has been outsourced to citizen science projects, such as Snapshot Serengeti (Swanson et al. 2015), or have turned to computer-assisted image analysis (Weinstein 2017).

Automated image processing algorithms for detecting, counting and identifying organisms in natural environments from camera trap images, aerial images or video streams have grown tremendously in the last decade (Weinstein 2017). A particular focus has been directed towards species-specific recognition algorithms (Weinstein 2017) based on unique species characteristics, such as shape, coloration or patterns (Bolger, Morrison, Vance, Lee & Farid 2012; Yu et al. 2013; Lhoest, Linchant, Quevauvillers, Vermeulen & Lejeune 2015). However, specific issues may arise from each study. For example, the Scale-Invariant Feature Transform (SIFT) algorithm (Lowe 2004), which recognizes distinctive animal features (spots, stripes, scales, or specific shapes such as fin or ear edges) on images (Bolger et al. 2012), is less efficient with highly flexible and deformable animals (e.g., foxes or cats) without any particular coat pattern (Yu et al. 2013). In general, deformable animals, which may exhibit different poses and shapes depending on the vision angle, are very difficult to detect against the background (Parkhi, Vedaldi, Jawahar & Zisserman 2011). In addition, automatic species identification algorithms are usually developed starting from selected images where the presence of an animal has already been validated by hand (Yu et al. 2013; Gomez Villa, Salazar & Vargas 2017), but automatic segmentation algorithms that allow the detection of animal presence/absence in images are still rare (Gomez Villa et al. 2017) or are case-specific. For example, AnimalFinder, which detects animals as large areas with low line density based on a canny edge detector, works well with medium-to large-bodied animals (such as white-tailed deer *Odocoileus virginianus*), but is less performant with short-statured or patterned species (Price Tack et al. 2016). More complex computer vision methods, such as deep learning algorithms, require a large training dataset where images have already been manually-labeled as containing or not an animal (Norouzzadeh et al. 2017). Partly to address this problem, the Where’s the Bear system uses stock Google Images to retrieve images of the target species and overlay them on empty, background images from the study site to construct a training set. The neural network training and subsequent image classification uses Google TensorFlow to recognize objects in images (Elias, Golubovic, Krintz & Wolski 2017).

For finding moving objects within images, a common technique in computer vision is background subtraction. In a sequence of images, it consists of subtracting the static background from the current image to detect its foreground (moving) objects. Having been mainly developed for video surveillance of buildings or streets, the existing background subtraction algorithms process video streams and need a relatively long sequence of frames to perform well (Ren, Han & He 2013). On the contrary, camera traps usually create short sequences of images at a low frame rate (usually 2-10 images at 1-2 frame/s) taken after a triggering event (Meek & Pittet 2012). Sequences of image files may be converted into a video file in order to use software that detect motion in videos (e.g., MotionMeerkat, Weinstein 2015), but these methods were not optimized for camera trap images (Swinnen, Reijniers, Breno & Leirs 2014). There is thus a clear need to develop background subtraction algorithms able to work from short sequences of images taken at a low frame rate, which is a typical output of camera trap studies.

Here we describe FoxMask, a new tool able to detect animal presence/absence in short sequences of camera trap images. FoxMask was primarily developed to analyze images taken on arctic fox *Vulpes lagopus* dens on Bylot Island (Nunavut, Canada). These images result from an intensive camera trapping study aiming at assessing fox presence and litter size from mid-May to early August. Arctic foxes in the tundra are challenging for automated image processing since they do not present any striking coat pattern or shape, are often partially hidden in the vegetation, and stay relatively low on the ground, especially as pups. Our goal was to reduce the amount of work related to ecological image processing by eliminating the need to inspect images that do not contain animals. We evaluated the performance of FoxMask under different conditions of natural background and lighting. The FoxMask program, detailed documentation and installation instructions are available at https://github.com/edevost/foxmask.

## FOXMASK

FoxMask is written in *Python* and *C++* and can be downloaded as an enclosed Ubuntu 16 Vagrant box. This box can be used for rapidly testing FoxMask and for development purposes, on any platform (e.g. Windows, MacOS, Linux). The box virtualizes a Linux guest operating system, from which the FoxMask software can be executed (option “Installation through a virtual machine”). Users can also use an automated installation script to install the software on their own Linux OS (option “Standalone installation on Linux”). FoxMask detects the presence of moving objects on images using background estimation and foreground segmentation algorithms. We refer to an *image sequence* when considering a group of consecutive images collected from one camera and to an *image series* when considering all the image sequences together. FoxMask is basically a wrapper of two open source algorithms developed by Reddy et al. (2011; 2013) that were further modified to process camera trap sequences. These algorithms were chosen for their efficiency to detect moving objects on short image sequences.

The execution of the FoxMask algorithm can be described by six major steps (Fig.1). (1) The metadata of each image are extracted and all images are sorted (increasing order) upon their time of creation. (2) Based on the sorted images list, sequences of consecutive images are created. For images to be considered as part of the same sequence, the time interval between each consecutive image has to be equal or less than the value entered in the user-defined *maxgap* parameter. Since our cameras were set to take five images after each trigger (see Dataset section below), for each image series, we chose a value of *maxgap* that would allow the selection of at least five images per sequence. The *maxgap* values used varied between 5 and 10 sec, and this resulted in sequences of 5-30 images (30 is the maximum number of images that FoxMask will consider as a sequence). (3) For each sequence, the static background is estimated (e.g. pixels that are considered statistically identical on all images are flagged as background; Fig. 2). (4) The detected static background is then fed to the foreground segmentation algorithm, where all images are subtracted from the reconstituted background. Following this operation, binary black and white masks are created and written to disk, with foreground objects (suspected moving ones) appearing in solid white, and the static background as solid black (Fig. 3). (5) Masks are then analyzed one by one and the area of each detected foreground object is calculated. This area constitutes a good parameter for distinguishing between animals and other moving objects often corresponding to background “noise” (for example, vegetation moved by the wind). The *minimum size threshold* (*minsize* parameter) represents the algorithm’s sensitivity to movement and should be set slightly smaller than the smallest animal targeted. (6) The final step consists in keeping only the images where at least one of the suspected moving objects has a size greater than the *minsize* set by the user. These retained images are those where at least one animal is present.

**Figure 1.**
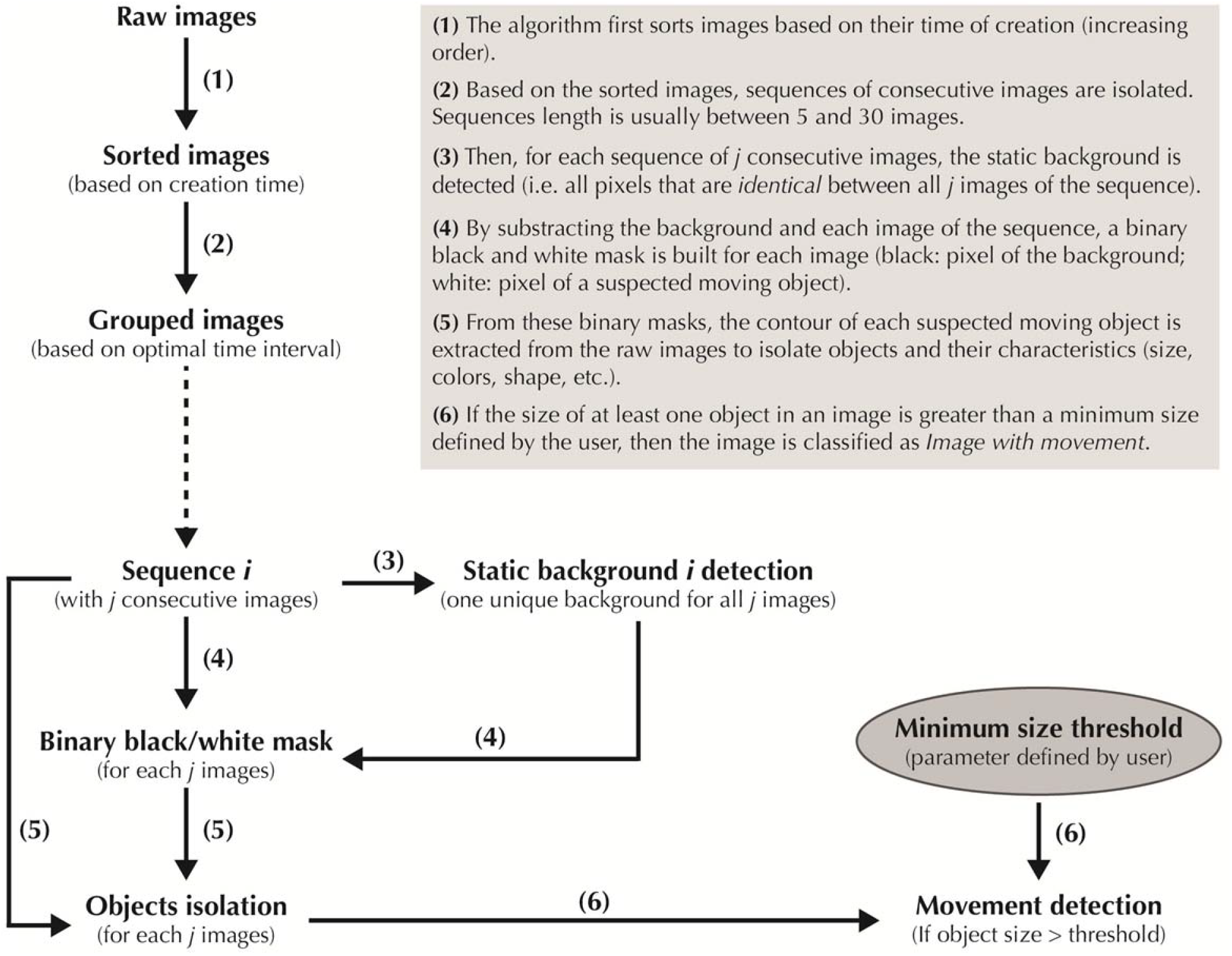
Summarized flowchart of data processing by the FoxMask algorithm.

**Figure 2.**
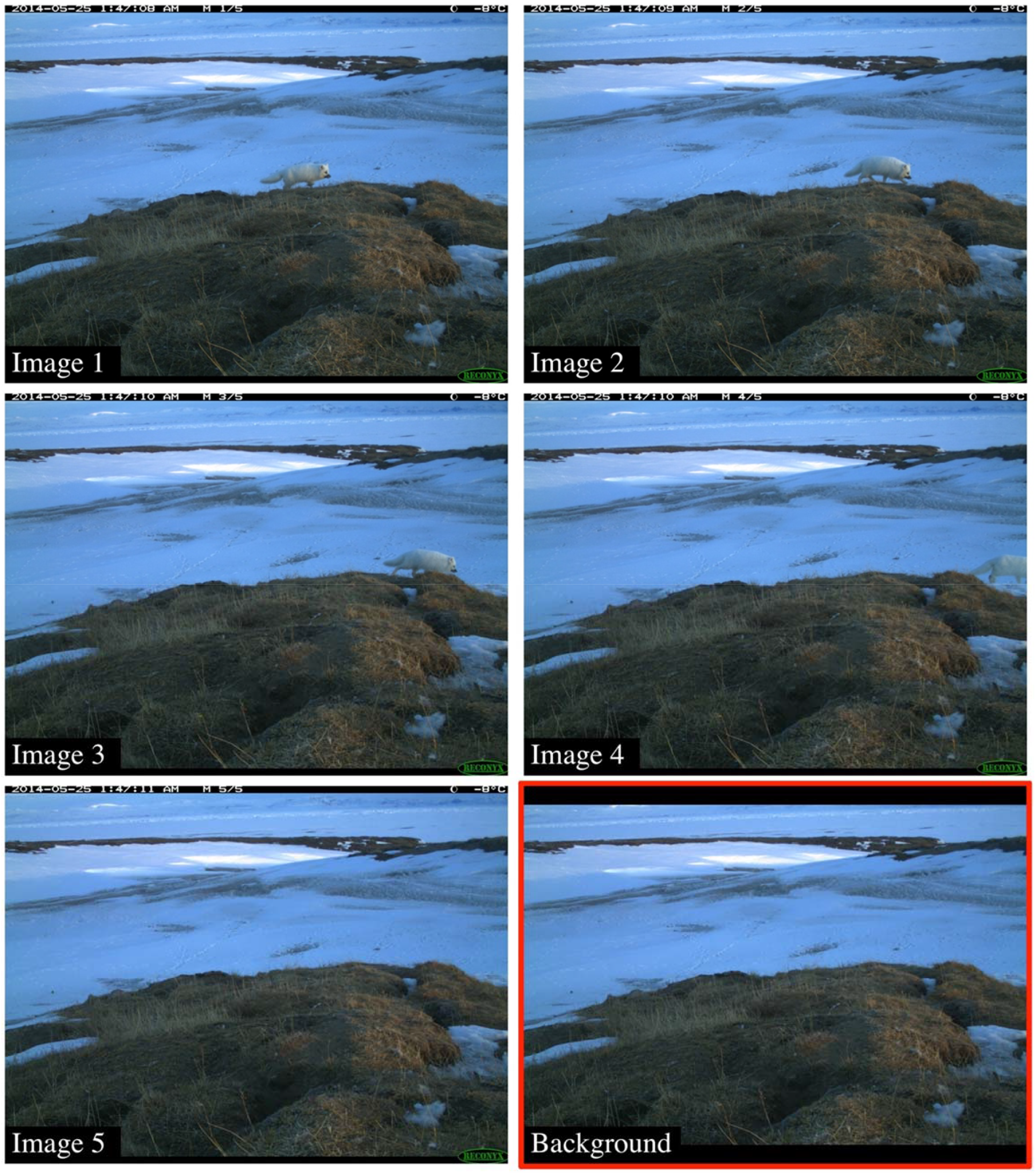
Example of the extracted static background (bottom-right) for a sequence of five consecutive images.

**Figure 3.**
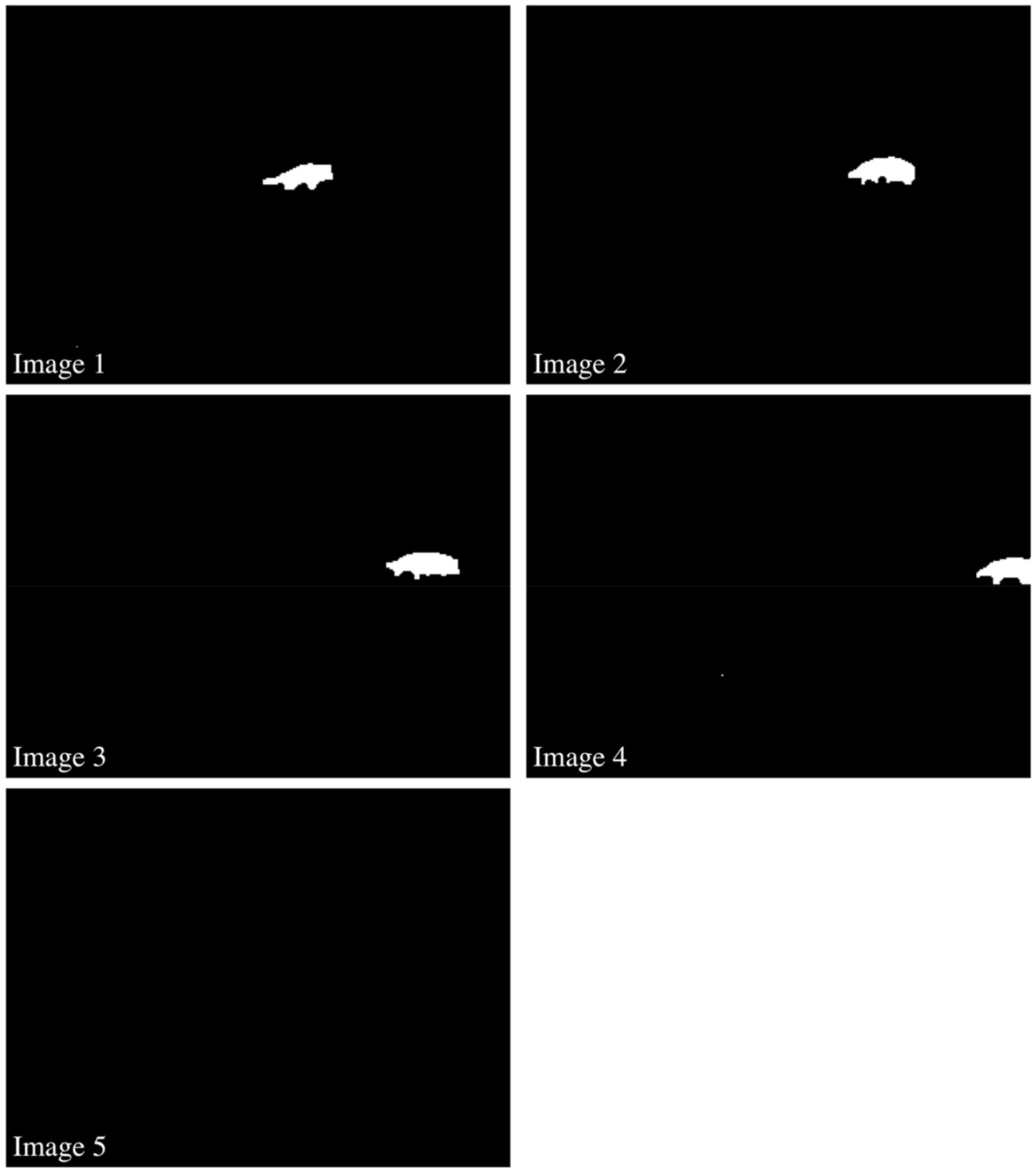
Example of five black and white masks obtained after the subtraction of the static background of the five original images. Pixels of suspected moving objects appear in white while pixels of static background appear in black.

For each folder of analyzed images, FoxMask provides (1) one table listing the names of all processed images and, for each image, a binary code indicating if at least one animal was detected or not (0: “No animal”; 1: “Animal”), and the minimum size threshold chosen, (2) a new subfolder containing only the images in the image series where an animal was detected.

## SENSITIVITY AND PERFORMANCE ANALYSES

### Dataset

We tested FoxMask on a dataset of images obtained from a camera trap monitoring of arctic fox dens on Bylot Island (73°N, 80°W). We used infrared automatic color cameras (RapidFire Professional PC85 and HyperFire PC800; Reconyx, Holmen, WI, USA) programmed to take a sequence of five photographs without time interval (*RapidFire* mode) each time a movement was detected. We tested FoxMask on 14,500 images that had been manually checked for animal presence or absence. “Animal” images could contain arctic foxes (adults and pups) or occasionally other species, such as red foxes *V. vulpes*, arctic hares *Lepus arcticus*, Greater snow geese *Chen caerulescens atlantica* and human observers. “No animal” images (also referred to as “empty images”) could either result from false triggers or the animal moving outside the camera field of view after a trigger.

### Sensitivity analysis

To examine the influence of the user-defined *minsize* on the classification of images (animal presence or absence), we used three image series of 500 consecutive images representing three scenarios. The first series had animals in 98.2% of images, the second series had animals on 52.0% of the images and the third series had no animal at all. We varied the values of the *minimum size threshold* from 0 to 5000 pixels (3000 pixels cover 1% of a picture). A low threshold would cause the algorithm to keep more images with background noise (small movements), while a large threshold would discard many images containing animals.

### Performance analysis

To evaluate the performance of FoxMask, we selected 26 image series of 500 consecutive images under different environmental conditions (Table 1). Based on results from the sensitivity analyses, we set the *minsize* at 200 pixels for all series. The proportion of images containing animals varied from 3.2 to 100 % (median = 86.1%). The performance of FoxMask was estimated as the proportion of images containing animals in the series correctly classified as “Animal”, the proportion of empty images in the series correctly classified as “No animal” and the proportion of all images in the series correctly classified in both categories. We visually examined all the images that the algorithm had not classified correctly to identify the potential case problems. If a detection error could have originated from a combination of two factors, both factors were counted as sources of error.

**Table 1.**
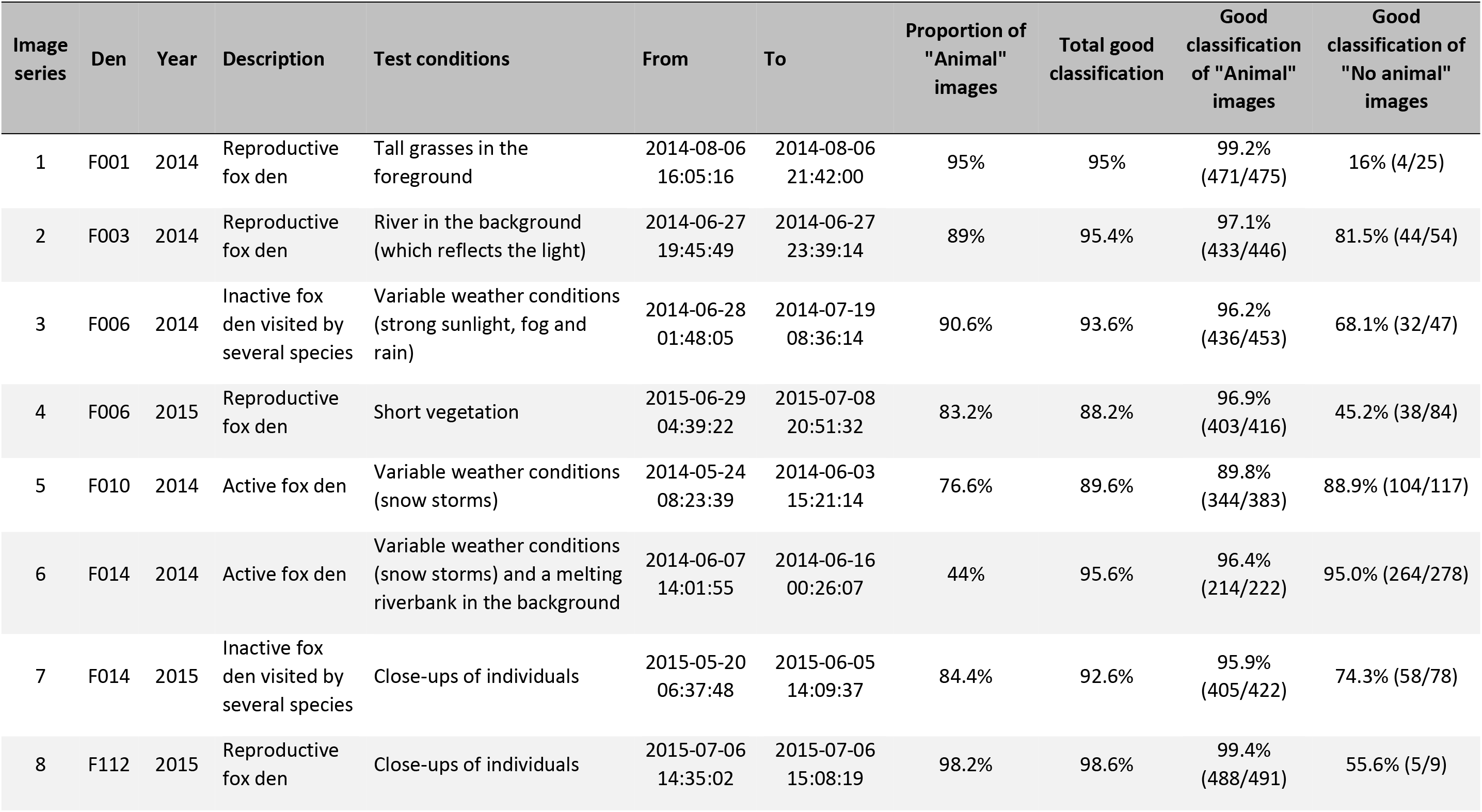
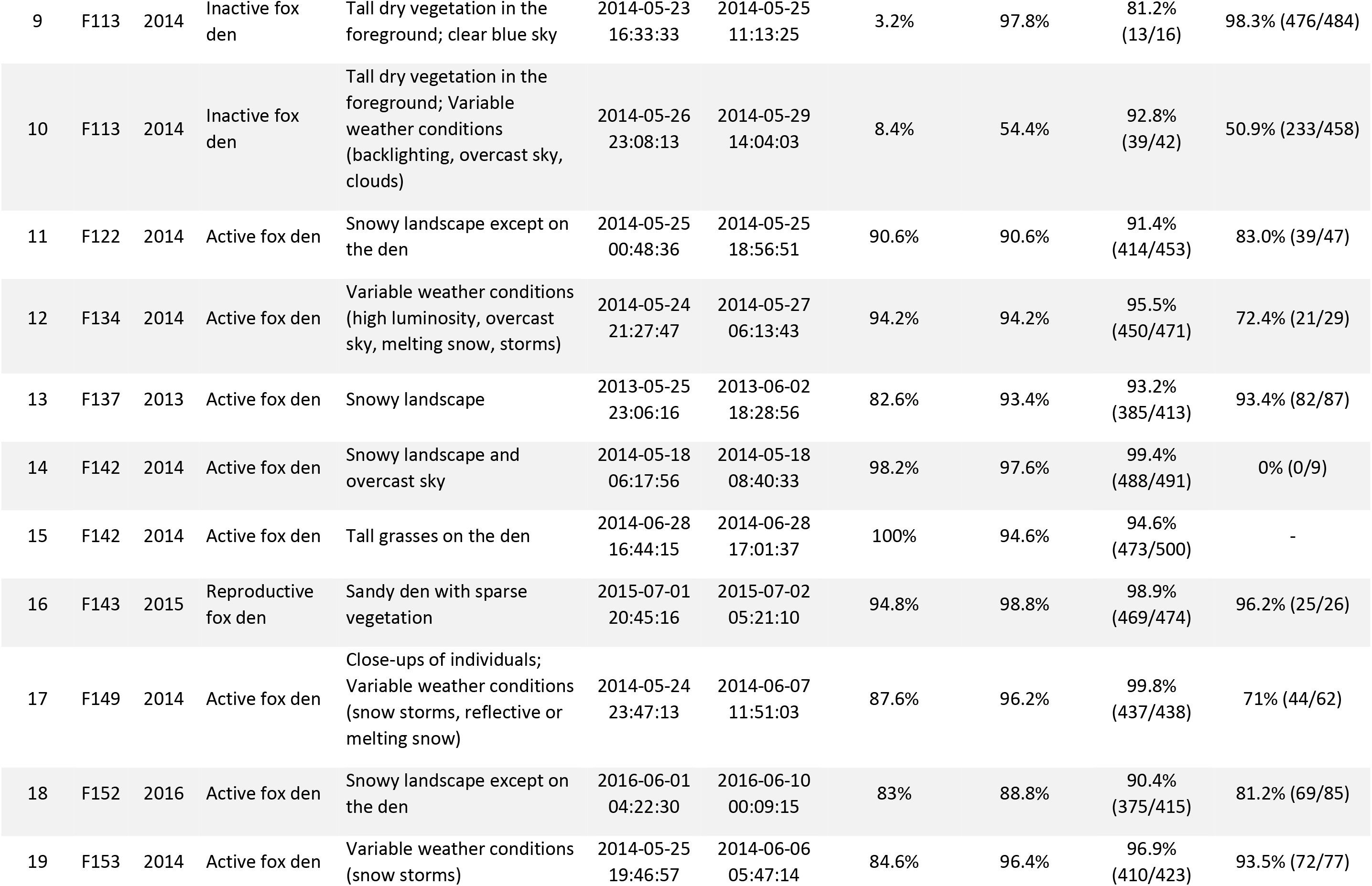
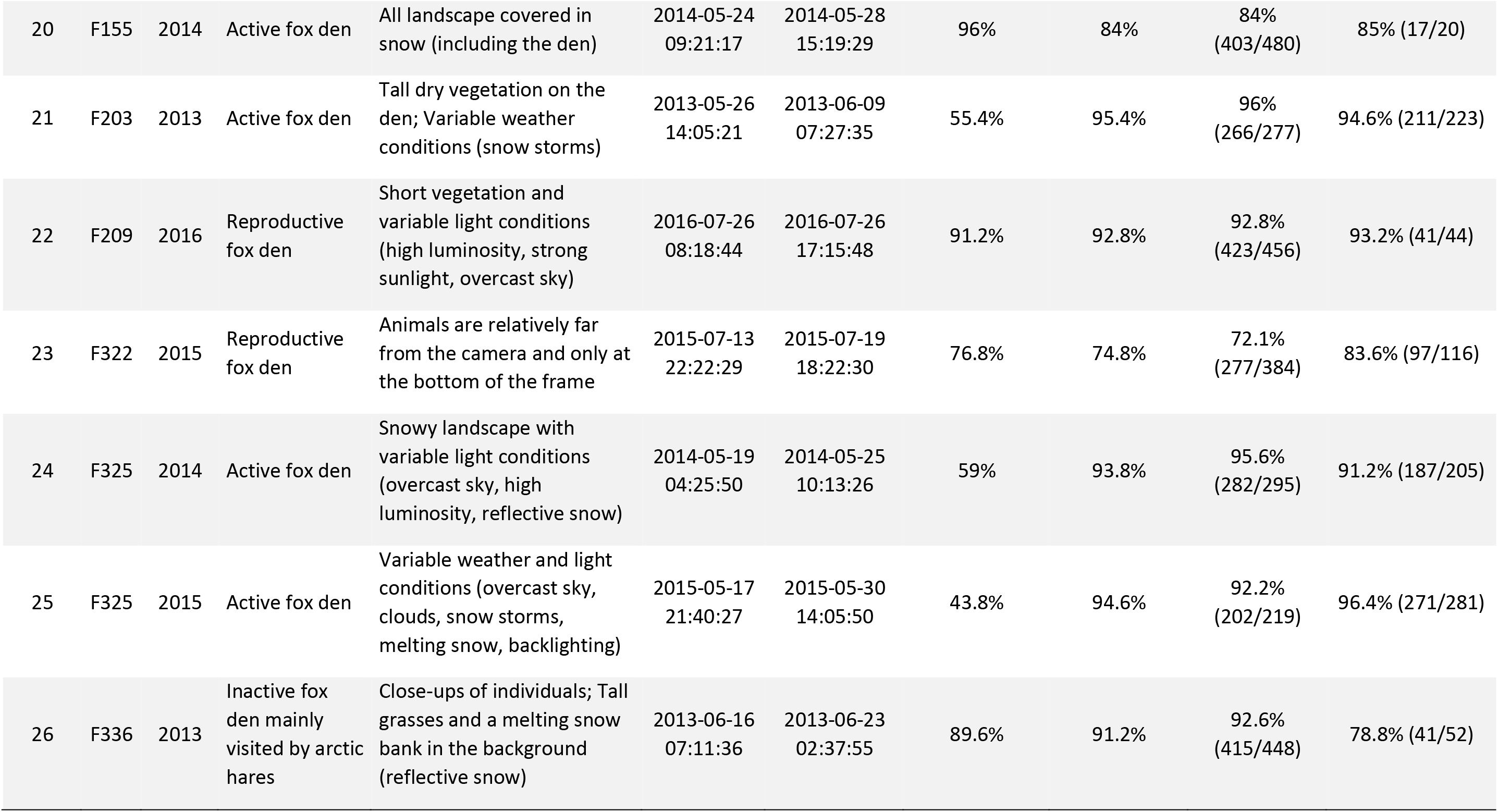
Description of the image series used to test FoxMask (n = 26) and results of the performance analysis. Reproductive fox dens were occupied by adults and pups, active fox dens were visited by adults (before pup emergence) and inactive fox dens were not regularly used by a fox pair, but may be occasionally visited by foxes and other species.

| Image series | Den | Year | Description | Test conditions | From | To | Proportion of "Animal" images | Total good classification | Good classification of "Animal" images | Good classification of "No animal" images |
| --- | --- | --- | --- | --- | --- | --- | --- | --- | --- | --- |
| 1 | F001 | 2014 | Reproductive fox den | Tall grasses in the foreground | 2014-08-06 16:05:16 | 2014-08-06 21:42:00 | 95% | 95% | 99.2% (471/475) | 16% (4/25) |
| 2 | F003 | 2014 | Reproductive fox den | River in the background (which reflects the light) | 2014-06-27 19:45:49 | 2014-06-27 23:39:14 | 89% | 95.4% | 97.1% (433/446) | 81.5% (44/54) |
| 3 | F006 | 2014 | Inactive fox den visited by several species | Variable weather conditions (strong sunlight, fog and rain) | 2014-06-28 01:48:05 | 2014-07-19 08:36:14 | 90.6% | 93.6% | 96.2% (436/453) | 68.1% (32/47) |
| 4 | F006 | 2015 | Reproductive fox den | Short vegetation | 2015-06-29 04:39:22 | 2015-07-08 20:51:32 | 83.2% | 88.2% | 96.9% (403/416) | 45.2% (38/84) |
| 5 | F010 | 2014 | Active fox den | Variable weather conditions (snow storms) | 2014-05-24 08:23:39 | 2014-06-03 15:21:14 | 76.6% | 89.6% | 89.8% (344/383) | 88.9% (104/117) |
| 6 | F014 | 2014 | Active fox den | Variable weather conditions (snow storms) and a melting riverbank in the background | 2014-06-07 14:01:55 | 2014-06-16 00:26:07 | 44% | 95.6% | 96.4% (214/222) | 95.0% (264/278) |
| 7 | F014 | 2015 | Inactive fox den visited by several species | Close-ups of individuals | 2015-05-20 06:37:48 | 2015-06-05 14:09:37 | 84.4% | 92.6% | 95.9% (405/422) | 74.3% (58/78) |
| 8 | F112 | 2015 | Reproductive fox den | Close-ups of individuals | 2015-07-06 14:35:02 | 2015-07-06 15:08:19 | 98.2% | 98.6% | 99.4% (488/491) | 55.6% (5/9) |
| 9 | F113 | 2014 | Inactive fox den | Tall dry vegetation in the foreground; clear blue sky | 2014-05-23 16:33:33 | 2014-05-25 11:13:25 | 3.2% | 97.8% | 81.2% (13/16) | 98.3% (476/484) |
| 10 | F113 | 2014 | Inactive fox den | Tall dry vegetation in the foreground; Variable weather conditions (backlighting, overcast sky, clouds) | 2014-05-26 23:08:13 | 2014-05-29 14:04:03 | 8.4% | 54.4% | 92.8% (39/42) | 50.9% (233/458) |
| 11 | F122 | 2014 | Active fox den | Snowy landscape except on the den | 2014-05-25 00:48:36 | 2014-05-25 18:56:51 | 90.6% | 90.6% | 91.4% (414/453) | 83.0% (39/47) |
| 12 | F134 | 2014 | Active fox den | Variable weather conditions (high luminosity, overcast sky, melting snow, storms) | 2014-05-24 21:27:47 | 2014-05-27 06:13:43 | 94.2% | 94.2% | 95.5% (450/471) | 72.4% (21/29) |
| 13 | F137 | 2013 | Active fox den | Snowy landscape | 2013-05-25 23:06:16 | 2013-06-02 18:28:56 | 82.6% | 93.4% | 93.2% (385/413) | 93.4% (82/87) |
| 14 | F142 | 2014 | Active fox den | Snowy landscape and overcast sky | 2014-05-18 06:17:56 | 2014-05-18 08:40:33 | 98.2% | 97.6% | 99.4% (488/491) | 0% (0/9) |
| 15 | F142 | 2014 | Active fox den | Tall grasses on the den | 2014-06-28 16:44:15 | 2014-06-28 17:01:37 | 100% | 94.6% | 94.6% (473/500) | - |
| 16 | F143 | 2015 | Reproductive fox den | Sandy den with sparse vegetation | 2015-07-01 20:45:16 | 2015-07-02 05:21:10 | 94.8% | 98.8% | 98.9% (469/474) | 96.2% (25/26) |
| 17 | F149 | 2014 | Active fox den | Close-ups of individuals; Variable weather conditions (snow storms, reflective or melting snow) | 2014-05-24 23:47:13 | 2014-06-07 11:51:03 | 87.6% | 96.2% | 99.8% (437/438) | 71% (44/62) |
| 18 | F152 | 2016 | Active fox den | Snowy landscape except on the den | 2016-06-01 04:22:30 | 2016-06-10 00:09:15 | 83% | 88.8% | 90.4% (375/415) | 81.2% (69/85) |
| 19 | F153 | 2014 | Active fox den | Variable weather conditions (snow storms) | 2014-05-25 19:46:57 | 2014-06-06 05:47:14 | 84.6% | 96.4% | 96.9% (410/423) | 93.5% (72/77) |
| 20 | F155 | 2014 | Active fox den | All landscape covered in snow (including the den) | 2014-05-24<br>09:21:17 | 2014-05-28<br>15:19:29 | 96% | 84% | 84%<br>(403/480) | 85% (17/20) |
| 21 | F203 | 2013 | Active fox den | Tall dry vegetation on the den; Variable weather conditions (snow storms) | 2013-05-26<br>14:05:21 | 2013-06-09<br>07:27:35 | 55.4% | 95.4% | 96%<br>(266/277) | 94.6% (211/223) |
| 22 | F209 | 2016 | Reproductive fox den | Short vegetation and variable light conditions (high luminosity, strong sunlight, overcast sky) | 2016-07-26<br>08:18:44 | 2016-07-26<br>17:15:48 | 91.2% | 92.8% | 92.8%<br>(423/456) | 93.2% (41/44) |
| 23 | F322 | 2015 | Reproductive fox den | Animals are relatively far from the camera and only at the bottom of the frame | 2015-07-13<br>22:22:29 | 2015-07-19<br>18:22:30 | 76.8% | 74.8% | 72.1%<br>(277/384) | 83.6% (97/116) |
| 24 | F325 | 2014 | Active fox den | Snowy landscape with variable light conditions (overcast sky, high luminosity, reflective snow) | 2014-05-19<br>04:25:50 | 2014-05-25<br>10:13:26 | 59% | 93.8% | 95.6%<br>(282/295) | 91.2% (187/205) |
| 25 | F325 | 2015 | Active fox den | Variable weather and light conditions (overcast sky, clouds, snow storms, melting snow, backlighting) | 2015-05-17<br>21:40:27 | 2015-05-30<br>14:05:50 | 43.8% | 94.6% | 92.2%<br>(202/219) | 96.4% (271/281) |
| 26 | F336 | 2013 | Inactive fox den mainly visited by arctic hares | Close-ups of individuals; Tall grasses and a melting snow bank in the background (reflective snow) | 2013-06-16<br>07:11:36 | 2013-06-23<br>02:37:55 | 89.6% | 91.2% | 92.6%<br>(415/448) | 78.8% (41/52) |

## RESULTS

On average, FoxMask analyses a folder containing 500 images in 20 min (Ubuntu 14.0 Box virtualized on an Intel Core i7 CPU with 2.9 GHz and 8 GB RAM).

### Sensitivity analysis

As expected, increasing the *minsize* resulted in FoxMask filtering out more empty images but also filtering in less images containing animals (Fig. 4a and 4b). Using an image series displaying a landscape in our study area but no images containing an animal, we were able to visually estimate the threshold for “typical” background noise. As seen in Fig. 4c, an asymptote starts to be reached at around 250 pixels, which corresponds to 95% of the empty images correctly classified as “No animal”. Based on the results on Fig. 4, we chose a threshold of 200 pixels that maximizes the correct classification of “Animal” images (over the correct classification of empty images) to conduct the performance analysis. For both series having “Animal” images, a value of 200 pixels results in approximatively 80% of good classification of all images. Although the algorithm was not able to classify all images correctly, this analysis showed it was possible to obtain a good trade-off between maximizing the elimination of empty images while minimizing the loss of “Animal” images in a series.

**Figure 4.**
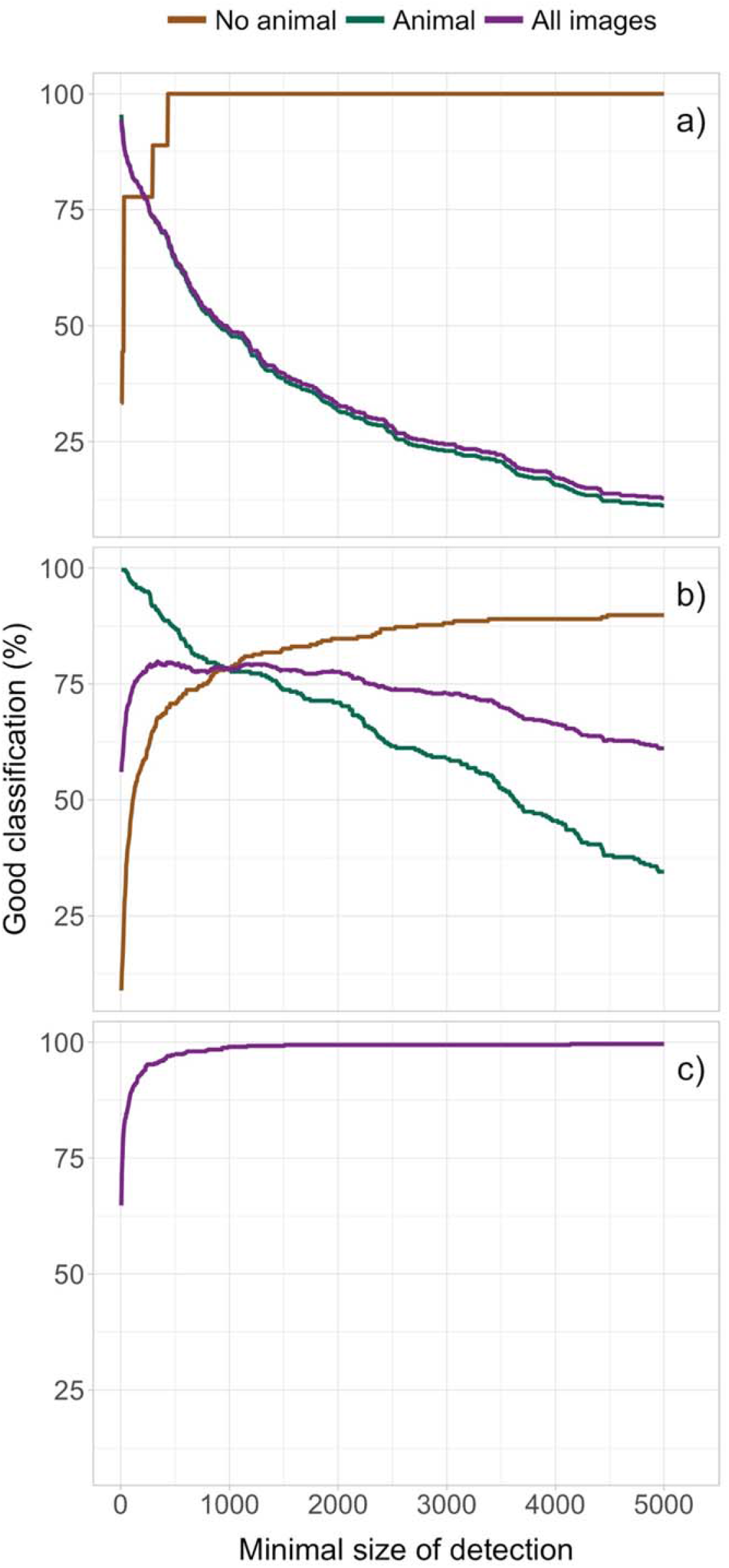
Impact of the user-defined minimum size threshold (3000 pixels cover 1% of an image), which controls the sensitivity of FoxMask to movement, on the proportion of images in the series correctly classified (purple line), as well as the proportion of images containing an animal correctly classified as “Animal” (green line) and empty images (brown line) correctly classified as “No animal”. Series of 500 consecutive images had (a) 97.2% of “Animal” images, (b) 52.0% of “Animal” images and (c) no “Animal” images. Note that in (c) the brown line is hidden below the purple line.

### Performance analysis

The average proportion of images (± SD) in each series correctly classified as “No animal” and “Animal” was 91.5% (± 9.1%; range = 54.4-98.8%). However, when the two categories were taken separately, the proportion of images containing animals correctly classified as “Animal” increased to 93.4% (± 6.2%; range = 72.1-99.8%) whereas the proportion of empty images correctly classified as “No animal” decreased to 75.4% (± 24.9%; range = 0-98.3%). The series with the lowest score (54.4% of overall correct classification) corresponded to an inactive fox den with tall dry grass in the foreground (Table 1). Although most “Animal” images in this series were correctly classified (39/42), 50.9% (233/458) of empty images were misclassified as “Animal” due to the moving grass in the foreground. Overall, the results of the performance analysis indicated that FoxMask generated few false negatives but quite many false positives in these difficult conditions of image analysis (Fig. 5).

**Figure 5.**
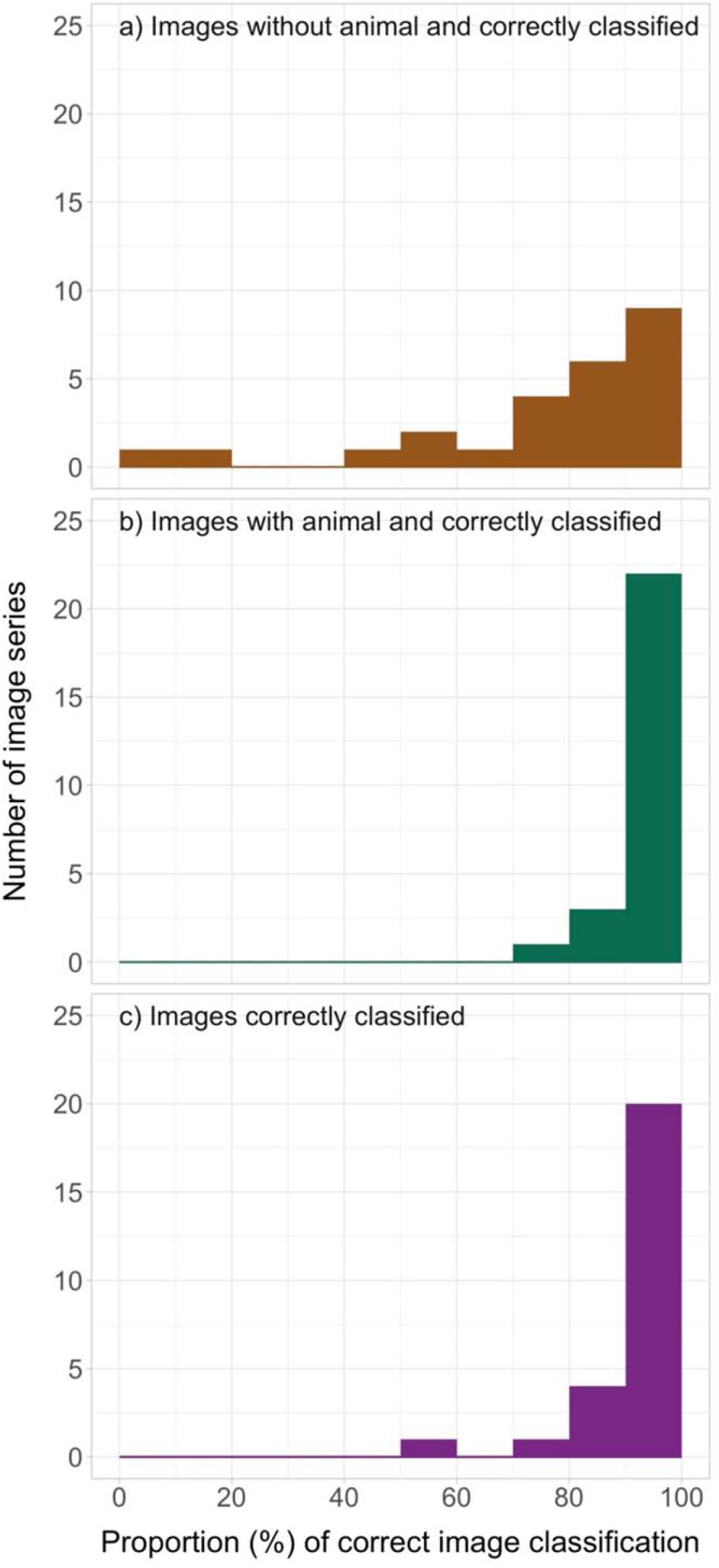
Number of image series (n = 26) belonging to each category of correct image classification after processing by FoxMask.

The post-hoc analysis of the images that were misclassified revealed the sources of error. The most frequent misclassifications arose from an animal being immobile or quasi immobile during a sequence of images (38.5% of 1146 cases). The static animal was sometimes considered as “background”, thus generating a false negative. At other times, when the animal moved out of the field of view at the end of the sequence of images, the landscape that was previously hidden was wrongly considered as a moving object, thus generating a false positive. Other sources of error, in decreasing order of frequency, were (1) false positive generated by foreground vegetation moved by the wind (22.2% of cases), (2) false negative generated by largely hidden animal (moving out of field of view, entering den, located behind obstacle, etc.) (11.4%), (3) false positive generated by a shadowed area (e.g., burrow entrance) interpreted as a moving object because of its changing darkness level (8.8%), (4) false negative generated by excessive distance of animal from camera (7.2%), (5) false positive due to the shadow of an animal appearing on the image while the individual was located out of the field of view (2.7%), (6) false positive generated by unsteady camera moving in the wind or abrupt change of ambient luminosity producing shifting background (3%). A few other sources of misclassifications were detected, such as a white arctic fox being too cryptic against the snowy background (false negative), an animal leaving deep footprints in the snow or moving patches of vegetation before leaving (false positive on subsequent images), or large snowflakes or heavy raindrops generating false positives (2.7%). We could not determine the source of misclassification in 3.1% of errors.

## DISCUSSION

With camera trapping having considerably facilitated field data collection, easy tools for automatic image analyses are needed in order to assist ecologists with data extraction. FoxMask reduces the workload during image analysis by automatically detecting animal presence/absence in camera trap images. Below we discuss the main advances in camera trap image classification brought by FoxMask, summarize our parameter recommendations, and highlight directions for future progress.

One difficulty faced by algorithm developers planning to use background detection to automate image analysis tasks in an ecological context is that natural landscapes (in contrast with parking lots, airport halls or jewelry stores) provide constantly changing backgrounds to moving objects. FoxMask can work with small sequences (5-30 consecutive images) to perform background subtraction, thereby allowing the static background to be constantly updated. This feature is essential when cameras stay in the field for a long time under changing weather, vegetation and luminosity. Although we mainly tested FoxMask with images of arctic foxes, it detects motion in general and can be parameterized for any species.

Our performance analysis indicated that the proportion of images correctly classified by FoxMask was very high (> 90% on average) and the algorithm worked well under various environmental conditions. Minimizing the number of images requiring a manual review can considerably reduce the time and costs of a camera trap monitoring program and speed up the data analysis. Depending on the parameter settings, FoxMask may however miss some images containing animals. It is therefore important to consider the trade-offs between the time saving benefits of the software, and its actual performance. An acceptable trade-off may be reached depending on the objectives of the research project. In general, if manually processing all images is the main workflow bottleneck in a research program, and analysis automation allows an increasing number of cameras to be deployed in the field, then false negatives are easily compensated by increased sample sizes of collected images.

We recommend determining the *minsize* parameter through a sensitivity analysis conducted on a sample of 100-500 images showcasing a typical background and the target species. Once the optimal threshold has been determined for this sample, if the conditions are relatively similar between cameras, this setting can be used to automatically batch-process multiple folders. Otherwise, if conditions vary considerably between cameras, we recommend choosing a *minsize* specifically for each camera. To maximize the performance of FoxMask, when placing cameras in the field, we recommend avoiding having vegetation (i.e., tall grasses or branches) and den holes or dark cracks in the foreground. We found that FoxMask failed to detect correctly animals that are standing relatively still. The algorithm may not be suited for target organisms that remain static for extended periods of time, such as an incubating bird.

Following the classification by FoxMask, the images flagged as “Animal” may then be analyzed using existing species identification algorithms (e.g., Yu et al. 2013) or an algorithm specifically designed by users. In our working case, the color of the objects could help identify adult foxes from pups since adults have either a white winter coat or a brownish-blond summer coat, while young foxes have a grayish fur. Another future step of FoxMask will aim to include a counting algorithm to estimate the number of individuals detected on images. Computer-assisted image analysis will most likely continue to improve in the next years and allow ecologists to increase their ability to collect meaningful biological data with a minimal time investment.

## ACKNOWLEDGEMENTS

We thank our many field assistants and students who helped collect the data. This project was supported by Canada Foundation for Innovation, Canada Research Chairs Program, Fonds de recherche du Québec - Nature et Technologies (FRQNT), International Polar Year program, Kenneth M Molson Foundation, Mittimatalik Hunters and Trappers Organization, Natural Sciences and Engineering Research Council of Canada (NSERC), Network of Centers of Excellence of Canada ArcticNet, Northern Scientific Training Program (Polar Knowledge Canada), Nunavut Wildlife Management Board, Parks Canada Agency, Polar Continental Shelf Program (Natural Resources Canada), Université du Québec à Rimouski (UQAR).

## AUTHORS’ CONTRIBUTIONS

DB and ED conceived the project; ED and NC developed the software, with input from SL; NC and SL analyzed the data and tested the software; SL and DB led the writing of the manuscript. All authors contributed critically to the drafts and gave final approval for publication.

## Notes

https://github.com/edevost/foxmask

